# Experimental loss of generalist plants reveals alterations in plant-pollinator interactions and a constrained flexibility of foraging

**DOI:** 10.1101/279430

**Authors:** Paolo Biella, Asma Akter, Jeff Ollerton, Sam Tarrant, Štěpán Janeček, Jana Jersáková, Jan Klečka

**Author notes:** Author for correspondence: Paolo Biella.

## Abstract

Species extinctions undermine ecosystem functioning, with the loss of a small subset of functionally important species having a disproportionate impact. However, little is known about the effects of species loss on plant-pollinator interactions. We addressed this issue in a field experiment by removing the plant species most frequently visited by insects, then measuring the impact of plant removal on flower visitation, pollinator effectiveness and insect foraging in several sites. Our results show that total visitation decreased exponentially after removing 1-4 most visited plants, suggesting that these plants facilitate others by maintaining high flower visitor abundances. Nevertheless, we found large variation in changes of visitation among plant species. Plant traits mediated the effect of removal on flower visitation; while visitation of plants which had smaller inflorescences and more sugar per flower increased after removal, foraging of flower visitors was constrained by not switching between flower shapes. Moreover, pollinator effectiveness fluctuated but was not directly linked to changes of flower visitation. In conclusion, it seems that the loss of generalist plants alters plant-pollinator interactions by decreasing pollinator abundance with implications for pollination and insect foraging. Therefore, generalist plants have high conservation value because they sustain the complex pattern of plant-pollinator interactions.

## Introduction

Overall community-level dynamics and ecosystem services are often disproportionately affected by a subset of the local species pool 1,2. This core of functionally important species contains, among others, generalists, which play a dominant ecological role within the community as they are often among the most abundant species3,4 and interact with a majority of other species5, mostly by complex facilitation-competition interactions6.

Generalist plants offer floral resources, mostly sugars from nectar and proteins from pollens, to a wide spectrum of pollinators and thus help to sustain those pollinators’ populations7. Pollinators of generalist flowers forage both as specialists and as generalists, creating a nested pattern of interactions in the entire plant-pollinator network, which, it has been argued, increases the robustness of the whole community8. Furthermore, visitation by a large number of different pollinators increases the chances of having the pollen dispersed9. In turn, when generalist pollinators are foraging in a patch, they can collect resources from a wide range of plants. This strategy could provide nutritional benefits from multiple sources10,11, but may also lead to pollination of several plant species because it is believed that generalist pollinators are visiting only a few plant species during each foraging bout9. Therefore, conservation of abundant generalists may be important, because their persistence can sustain most of the complexity of interactions taking place in a community12.

Despite their functional importance, little is known about the effects of local extinction of generalists. The consequences of species loss have traditionally been investigated by removing species at random. However, in nature this is rarely the case as species do not go extinct randomly, but rather non-random losses are the rule13–15. In the context of plant-pollinator interactions, the loss of plants or pollinators and its consequent effect on the interacting assemblages has been studied mostly by simulation models16,17. Experimental tests in the field have started only recently and have been so far limited to relatively small manipulations, for instance removing either one invasive plant species18,19 or a single native plant species20. Other studies have excluded one species or a small set of pollinators21,22. Furthermore, only simulation-based studies have tested the effects of excluding several species sequentially17,23, while the few field experiments removed only one species and led to disparate results. For instance, removal of one species of bumblebee led only to small differences in the interaction networks after manipulation21, while a study removing one invasive plant species found a pronounced effect of removal18. In fact, removing either a pollinator or a plant has different implications. Specifically, by removing a generalist pollinator, more resources will be available for the other pollinators; conversely, the removal of an abundant plant induces an immediate reduction of resources available to the pollinator guild. However, the consequences of the loss of multiple species for plant-pollinator interactions remains an open question.

In this study we experimentally tested the effects of the removal of generalist plant species on plant-pollinator interactions. We focused on pollinators as a guild rather than on individual pollinator species by investigating the effects of the removal of the main flower resources on visitation, nectar consumption, effectiveness of pollination and shifting resource use by the pollinator guild. The experiments comprised two parts. Firstly, a pilot project in the United Kingdom experimentally manipulated a plant-pollinator community by removal of the most generalist plant (from here on called the “pilot study”). This provided a proof-of-concept and suggested ways in which the pollinator assemblage might react to such perturbation. Secondly, in the Czech Republic, we conducted a larger experiment, where we sequentially removed four of the most visited plant species over a short time period to determine how the plant-pollinator interactions would respond to a more profound loss of resources (from here on called the “sequential removal” experiment).

The aim of our experiments was to test whether removal of generalist plants (1) led to a decrease in the overall visitation i.e., the abundance of pollinators in the sites; (2) caused species-specific changes of the visitation of individual plant species, which could be explained by shifts of pollinators driven by traits of the plants, and (3) whether pollination effectiveness (determined by the number of pollen tubes grown in the pistils after visitation) and the amount of used nectar resources (“standing crop”) changed as a consequence of plant removal.

We hypothesised that the pollinator guild could respond to the removal of major floral resources in three ways: (a) they could shift food source and distribute evenly on the remaining plant species; (b) they could shift preferably towards a subset of the plants, possibly on the basis of plant traits; (c) they could stop foraging at the site (or emigrate) due to the lack of the main food source. The scenario (a) might be expected if pollinators are generalists able to use any resources. In addition we hypothesised that under scenario (a) and (b) plant reproduction, measured as number of pollen tubes, would increase, whereas under scenario (c) it would decrease.

## Methods

### Pilot study

The pilot study was conducted in the East Midlands of the UK, in Northampton, during summer 2008 on the 3 ha Quarry Field (52°16’12 N, 0°52’46W), part of the Bradlaugh Fields site, a network of parkland and Local Nature Reserves (data in S1 Table).

Surveys of insect visitation were undertaken at two stages: (i) before flower removal in order to determine the plant-pollinator interactions prior to the experiment; (ii) in the days following the removal of inflorescences of the plants species with the highest visitation, i.e. the total abundance of flower visitors on the plant. Inflorescences of the target plant (*Knautia arvensis*) were removed from the whole surface of the entire site. To indicate which plant to be removed, flower visitor abundance was used because it would not be practical to wait for species-level identification of flower visitors needed to decide which was the most generalised plant. We later confirmed that the plant with the highest abundance of flower visitors was also the one with the highest species richness (see Results). During the same period, the nearby Scrub Field, which hosts a vegetation similar to the treated site, was also surveyed two times using the same techniques, as a comparison control site to the Quarry Field. Insect flower visitation surveys were undertaken four times at each of the two stages of the experiment (i.e., before and after removal) between 1pm and 4pm on days which were warm and sunny with little or no wind. Surveys lasted 30 minutes and all flower visiting insects seen to be feeding within the flowers were captured along a 2 metre wide belt and within 2 metres in front of the surveyor. The sampling followed a widening spiral from a randomly determined point at a standard pace of 10 m per minute (which makes each survey of an area of approximately 600 m^2^). This method allowed for a large area to be surveyed owing to the relatively low pollinator density where floral resources are patchy and sparsely distributed. Insect specimens were identified either in the field or by experts. Sampling permission was obtained from The Wildlife Trust for Bedfordshire, Cambridgeshire, Northamptonshire and Peterborough.

### Sequential removal

The sequential removal experiment was performed in the vicinity of Český Krumlov (South Bohemia, Czech Republic) in the summer 2015. No sampling permits were required for this project in the Czech Republic, because no protected species were affected and the study was conducted on public land. Three experimental sites were chosen (Site 1: 48° 49′ 26.8′′ N, 14° 16′ 26.2′′ E, area ca. 1500 m^2^; Site 2: 48° 49′ 51.63′′ N 14° 17′ 34.12′′ E, ca. 1800 m^2^; Site 3: 48° 49’ 35.07’’ N 14° 18’ 8.2’’ E, ca. 1600 m^2^). The mean pairwise distance between the sites was 2.01 ± 0.95 Km. These experimental sites were small dry meadows surrounded by trees or bushes and with a dense forest on at least two edges. These were intended as physical barriers clearly separating the experimental sites from other grassland habitats in the surroundings. The entire surface of each experimental site was treated by removing the target plant species (see below).

An untreated control site was located at 48° 49′ 26.8′′ N, 14° 16′ 26.2′′ E, which consisted of habitat and plant community very similar to the treated sites. It was not feasible to pair each experimental site with a control site because of the lack of sufficiently similar sites in the vicinity of the experimental sites, so we opted for a single control accompanying the three experimental sites. An alternative design such as one where we would manipulate one half of a site and use the half as a control would violate the independence of treated or untreated plots given the flying ability of pollinators and the lack of separation of the plots by physical barriers or by distance.

The sampling of insects was based on walking six transects (10 m x 1 m) in a randomized order in each site between 9:00 and 17:00 hours. Transects were set up for sampling pollinators and to count plant abundances in order to account for heterogeneity in plant distribution within the sites. The size and number of transects was set proportionally to fit within the small size of the selected sites. All transects were usually sampled twice during each day. Sampling was postponed in the case of rain or strong wind. All insects found while visiting flowers were sampled by a handnet or a mouth aspirator. The floral abundance of plant species was recorded by counting the number of open flowers or compound inflorescences on each plant within each transect. These data were recorded during all stages of the experiment, so that changes in plant phenology across the experimental period would be recorded as changes in the number of flowers, which is the appropriate measure of plant abundance in the context of plant-pollinator interactions. It was not possible to collect further details on the flower sexual maturation stage (pollen presentation or stigma receptivity) because that usually requires destructive methods^25^, which could affect the flowering community in the sites, and it would also not be feasible to collect such data for the entire community given the large number of species present.

In the experimental sites, all flower visitors feeding on flowers were sampled for two days prior to any manipulation (hereafter the “Before” period). At the end of the sampling days, the captured specimens were counted for each plant and thus we were able to determine the plant species with the highest visitation, i.e. the total abundance of flower visitors on the plant. We call these highly visited plants “generalist”, although we did not evaluate the diversity of the visitors, in accordance with the literature (see ^20^) and the outcome of the pilot study (see Results). This most generalist plant species was then removed from the entire site by cutting all inflorescences in the entire site, as was done in the pilot study. We only removed flowers and left the stems otherwise intact so that the vegetation structure remained unaltered. Twenty four hours after the removal, the sites were sampled again over two days. After we had counted the abundances of the visitors for each plant, we determined the next plant species with the highest visitation, the inflorescences of which were then removed. The flower visitors were sampled again after another twenty four hours for two days. We repeated this procedure until the fourth plant species was removed, which was followed by the last sampling period. Throughout the experiment, we verified that inflorescences of the removed plants were still absent in the sites. Site 1 and 2 were sampled from 25th June to 12th of July, Site 3 was sampled from 2nd to 17th of July. The control site was sampled synchronously to each of the experimental sites (the same days and during the same hours), but no manipulation of the plant community was performed there (data in S2 Table). For each site, we analysed data on total flower visitation of all plant species by the entire guild of flower visiting insects, data on insect species identity from this experiment are yet not available. The sequence of the plant species removal for both pilot study and sequential removal is detailed in Table 1.

**Table 1.** Plant species removed during the experiments in the treated sites, with details on the plant family and the raw number of specimens found on that given plant.

| Treatment period | Species removed | Family | Number of specimens |
| --- | --- | --- | --- |
| Pilot study |  |  |  |
| Before removal | *Knautia arvensis* | Caprifoliaceae | 54 insects<br>(of 11 species) |
| Sequential removal experiment |  |  |  |
| Before removal |  |  |  |
| Site 1 | *Anthriscus sylvestris* | Apiaceae | 256 |
| Site 2 | *Veronica teucrium* | Plantaginaceae | 103 |
| Site 3 | *Aegopodium podagraria* | Apiaceae | 141 |
| 1 sp. removed |  |  |  |
| Site 1 | *Rubus caesius* | Rosaceae | 315 |
| Site 2 | *Agrimonia eupatoria* | Rosaceae | 96 |
| Site 3 | *Veronica teucrium* | Plantaginaceae | 26 |
| 2 spp. removed |  |  |  |
| Site 1 | *Knautia arvensis* | Caprifoliaceae | 83 |
| Site 2 | *Centaurea scabiosa* | Asteraceae | 96 |
| Site 3 | *Knautia arvensis* | Caprifoliaceae | 45 |
| 3 spp. removed |  |  |  |
| Site 1 | *Galium mollugo* | Rubiaceae | 39 |
| Site 2 | *Securigera varia* | Fabaceae | 26 |
| Site 3 | *Origanum vulgare* | Lamiaceae | 55 |

### Pistil collection and pollen tubes

Pollinator effectiveness was tested by counting pollen tubes in the pistils of flowering plants in two of our experimental sites. Growth of pollen tubes provides information about the effectiveness of the pollination service because it links pollen deposition and seed production^26^, although it may confound cross and self pollination. We collected pistils from on average 40 flowers of each sufficiently abundant flowering plant after each insect sampling period and preserved in formalin-acetic acid-alcohol (FAA) at room temperature. Flowers were collected outside (but nearby) the transects to avoid depletion of flower abundances in the transects. Later, in the laboratory, the flowers were dissected under the microscope and pistils were prepared for softening and staining, following the technique of ^27^. After softening the pistils in 4M NaOH, they were stained with 0.1% aniline blue in 0.1M K_2_HPO_4_ for 12 hours. Then, the pistils were washed and mounted in a drop of 50% glycerine on glass slides and covered with cover slips for observation under a fluorescence microscope. Pollen tubes were visible and counted for most of the species. However, in a few cases, pollen tubes were impossible to visualize properly. For these species the number of pollen grains on the stigmas were counted assuming that only germinated grains with tubes still attached would remain on the stigma after the preparation process. All processes were carried out at room temperature. After the observation, the edges of the cover slips were sealed with clear nail polish and stored at 4°C for future reference. Data were successfully obtained for 10 species because other plant species did not yield any countable pollen tubes (data in S3 Table).

### Nectar content of flowers and other functional traits

We determined the standing-crop of nectar in flowering plant species at two sites of the sequential removal experiment in order to assess the amount of unused floral resources and its changes after plant removal. To do so, we collected flowers in each site after the pollinator sampling on the same day (data in S4 Table). A 100μl Hamilton capillary syringe was used for the collection and for washing the nectar into distilled water. Flowers were selected randomly but outside the sampling transects in order to avoid impoverishment of flower resources in the transects; only flowers in full anthesis were sampled. Nectar samples were stored in a cool bag in the field and in a −20°C freezer in the laboratory. Sugar analysis of nectar was done using high performance anion exchange chromatography with pulsed amperometric detection using a Dionex ICS-3000 system and CarboPac PA1 analytical column. Nectar amount was expressed as milligrams of sugars per flower. Because the method is not sensitive when the amount of sugar is extremely low, several nectar samples from a known number of flowers (of a given species on a given day) were merged in one unique sample which was analysed as described above; afterwards, the amount of sugar per flower was calculated by dividing the quantity of sugar by the number of flowers included in the sample. Data were extracted from an average of 45 flowers per plant species.

Furthermore, we measured several functional traits for each plant species in the two sites. The daily production of nectar was measured with a similar methodology as for the standing crop but the only difference was that flowers were bagged for 24 hours before sampling nectar: several inflorescences were bagged in the morning and sampled the next day in the morning, yielding nectar data on average 45 single flowers per species. This is an appropriate method used for comparing the cumulative secretion of nectar over a standard amount of time which covers the entire daily rhythm in several different species. Other traits were: plant height (linear distance between the ground and the top of an inflorescence, measured on an average of 12 plants per species); inflorescence maximum size (the maximum dimension of the inflorescence, measured on an average of 10 flowers per species); dominant colour of the corolla (categorical variable: “white”, “blue”, “pink”, “yellow”); flower shape coded according to ^7^ as follows: bell- or funnel-shaped, dish- or bowl-shaped flowers, flag-shaped flowers, gullet-shaped flowers, head- or brush-shaped, tube-shaped flowers; unclear cases were checked with ^28^.

### Statistical analyses

All data were analysed by means of generalized linear mixed-effect models (GLMM) using the library *lme4* ^29^ in the R environment ^30^.

### Site level visitation analyses

For the sequential removal, the overall visitation was derived by calculating the sum of flower visitor abundances across plant species for each transect, separately at each sampling event. Thus, the number of flower-visiting insects per transect was the response variable, treatment was a categorical predictor describing the number of removed plants (“Before”, “1 sp.”, “2 spp.”, “3 spp.”, “4 spp.”). Transect identity nested within site identity was used as a random effect on the intercept and a Poisson distribution was used. We used plant visitation as an offset in the model to account for possible confounding effects caused by factors other than our experimental manipulation, e.g. phenology, decrease of insect abundance due to sampling, etc. Thus, the plant visitation in the control site at a given time (geometric mean across the control site’s transects) was included in the model as an offset. In the experimental sites, a multiple comparison tests were performed using the package *multcomp* with the function *glht*^31^ to compare a given treatment level with the preceding treatment level (e.g. “1 sp. removed” vs “Before”, “2 spp. removed” vs “1 sp. removed”, and so on) in order to test the significance of relative increases or decreases in pollinator abundances.

For the data from the pilot study, the visitation was analysed in a slightly different fashion because flower visitors were collected along a single transect and insect species were taxonomically identified. Thus in a GLMM with Poisson distribution the number of visits by individual flower visitor species at each plant species was used as a response, treatment was a predictor (“before” and “after” removal of *Knautia arvensis*). Species identity was used as a random factor on the intercept. The analysis included visitation in the control site as an offset for the same reasons as for the sequential removal (see above). Specifically, visitation by an insect species on a plant species in the control site was used as an offset for visitation in the same insect-plant combination in the experimental site. For those insect species that were not recorded in the control site, the mean visitation recorded on that plant species across all visiting insect species at a given experimental time was used. Data from the control site were also analysed in a similar way (but without an offset); i.e. we also tested whether plant visitation changed between the sampling periods in the control site. For the control site, changes in the visitation on the species that was removed from the treated site were explored with a similar GLMM model, but without plant species as the random intercept.

### Plant species level visitation analyses

To test the effect of treatment on each plant species in the sequential removal experiment, plant-level visitation (visiting insect abundance for each plant species at each sampled transect, including unvisited plants) was the response variable and treatment was a predictor variable of the removal events (“Before”, “1 sp.”, “2 spp.”, “3 spp.”, “4 spp.”). An offset of plant abundance measured as the number of flowers per transect was included, to account for possible variation of the amount of flowers during the experiment (changes in plant abundance or phenology), as suggested in^32^. Transect identity within site identity was used as a random intercept. Plant species identity was used as another random factor affecting the slope of the treatment; i.e. we assumed that different plants may respond to the removal treatment in a species-specific way. Poisson distribution was used in the GLMMs.

### Pollination effectiveness analyses

We tested whether the number of pollen tubes in pistils of each plant species, i.e. viable pollen grains deposited and successfully germinating, changed during the experiment in the sequential removal sites. The number of pollen tubes was used as a response variable, the number of removed plants (coded as a categorical variable: “Before”, “1 sp.”, “2 spp.”, “3 spp.”, “4 spp.”) and the plant’s mean visitation across each transect were used as predictors. Random slopes of predictors were included with site identity within plant species as a random factor; i.e. we assumed that the effect of the predictors varied between plant species and sites. The GLMM was fitted with a Poisson distribution.

### Standing crop of nectar analyses

To test the variation of standing crop of nectar, the amount of sugars per flower of each plant species was used as a response variable, the treatment of the removal events (“Before”, “1 sp.”, “2 spp.”, “3 spp.”, “4 spp.”) and the plants’ mean visitation across transects were used as predictors as in the analysis of pollen tubes (see above). We used site identity within plant species as a random factor affecting the intercept, i.e. assuming that the mean amount of sugar per flower varies between sites and plant species (including the effect of the random factors on the slope was not possible due to a lack of convergence of the model). Gamma distribution was used, and the link function was the natural logarithm.

### Traits analyses

To test if foraging during the treatment would be directed towards flowers being similar or dissimilar to the removed species, we assessed how visitation related to differences between plant-species’ traits and the removed plants’ traits in the two sites where traits were measured (data in S5 Table). To do that, the log-ratio difference between a given continuous plant trait value and the trait value of the removed plant species, *log(trait_sp. i_ / trait_removed sp._),* was calculated for each treatment level and used as a predictor variable. This measure of difference between quantitative values thus varies from negative values, through zero to positive values and has favourable statistical properties for analysis^33^. For categorical variables, a binomial variable was used, i.e. “same” as or “different” from the removed species. We aimed to compare visitation of individual plant species before and after each removal event (i.e. 1 sp. removed vs. before, 2 spp. removed vs. 1 sp. removed, etc.), so we reorganised the data for this analysis with the treatment coded simply as “before” and “after” removal. Plant traits used as predictors of visitation changes were flower colour, flower shape, inflorescence size, plant height and the daily production of sugars in nectar. We tested whether the change in visitation of individual plant species was affected by its traits by including interaction terms between the traits and the treatment variable. By including the treatment as predictor, the variation of visitation amount over the experimental time is included in the model. The number of flower of each plant species was included as an offset to account for possible variation of plant abundance across treatments. Transect identity within the site identity was used as a random intercept. Plant species identity was used as another random factor affecting the slope of the treatment effect. A Poisson distribution was used in the model. The control site was not included in this analysis because species were not removed in that site and thus it is not possible to calculate the trait similarities between the removed and remaining plant species.

## Results

In the pilot study, the surveys identified a total of 13 insect pollinated plant species in flower and 25 pollinating insect species in the experimental site. The plant with the highest pollinator abundance and also the highest visiting species richnes was *Knautia arvensis* (details in Table 1). Most other plants were visited by fewer than 20 insect individuals and less than 8 species in both experimental phases, except *Centaurea nigra* which was visited by 41 individuals of 11 species before removal and 86 individuals of 11 species after removal of *Knautia*.

In the sequential removal experiment, the sites 1, 2 and 3 were surveyed for a total of 31, 26 and 23 insect pollinated plant species, respectively. The amount of insect specimens and the sequence of removed plants is detailed in Table 1.

### Results from the pilot study

In the pilot study, visitation to plants changed only slightly after removing one species (Fig. 1). The treatment was not a significant predictor of visitation in the statistical model including an offset of the control’s visitation when compared with a model without treatment variable (**χ**^2^=0.194, df=1, p = 0.66).

**Figure 1.**
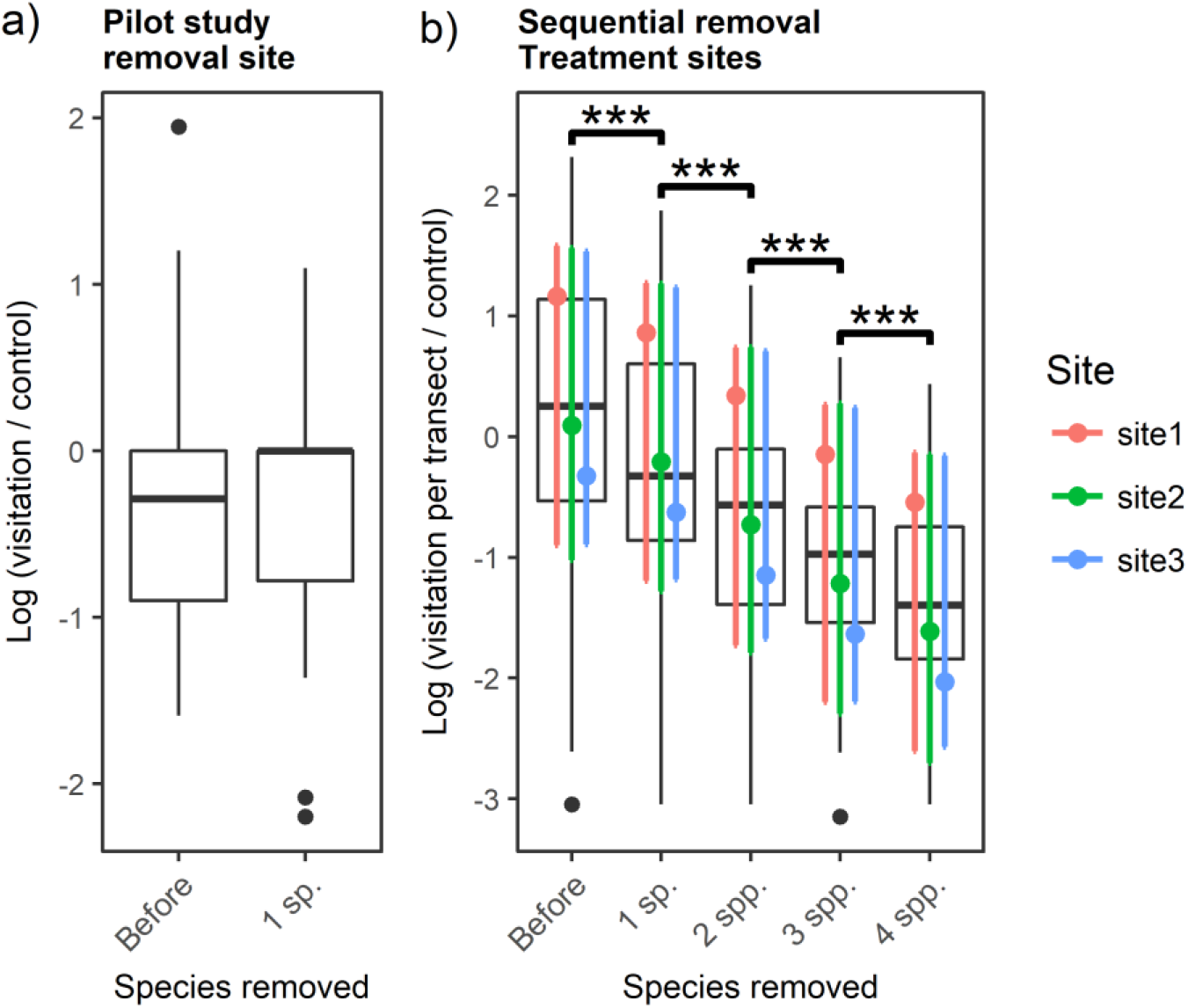
Overall visitation in both pilot study and sequential removal experiment, including offsets of control sites’ visitation. (a) Boxplots of raw data of the pilot study indicating the flower-visitor visitation (abundances) for each plant species; (b) Plot of the sequential removal experiment in where the boxplots indicate visitation across plant species of the same sampling event (i.e. a transect walk) and the estimated (modelled) means and confidence intervals are represented. All plots are on a logarithmic scale. When significant after multiple comparison test, it is indicated with the following codes: *** p<0.001, ** p<0.01, * p<0.05.

Visitation changed only slightly in the near control site, the difference was not statistically significant (**χ**^2^=0.85, df=1, p = 0.36) (Fig S1). In the control site, visitation to *Knautia arvensis* (the plant that was removed in the experimental site) did not change over the study period when compared with a model without treatment variable (**χ**^2^=0.852, df=1, p = 0.36).

### General pattern of visitation in the sequential removal experiment

In the sequential removal sites, visitation per transect decreased sharply after the removal of selected plants. The treatment was a significant predictor when compared with a model without treatment variable (**χ**^2^= 1605.3, df=4, p < 0.001), in the model including the visitation in the control site as an offset. The model without offset of the control site’s visitation yielded very similar results. Multiple-comparison test of treatment levels gave significant results in most cases (Table 2), with a similar decrease of total visitation after each stage of the removal experiment (Fig. 1).

**Table 2.**
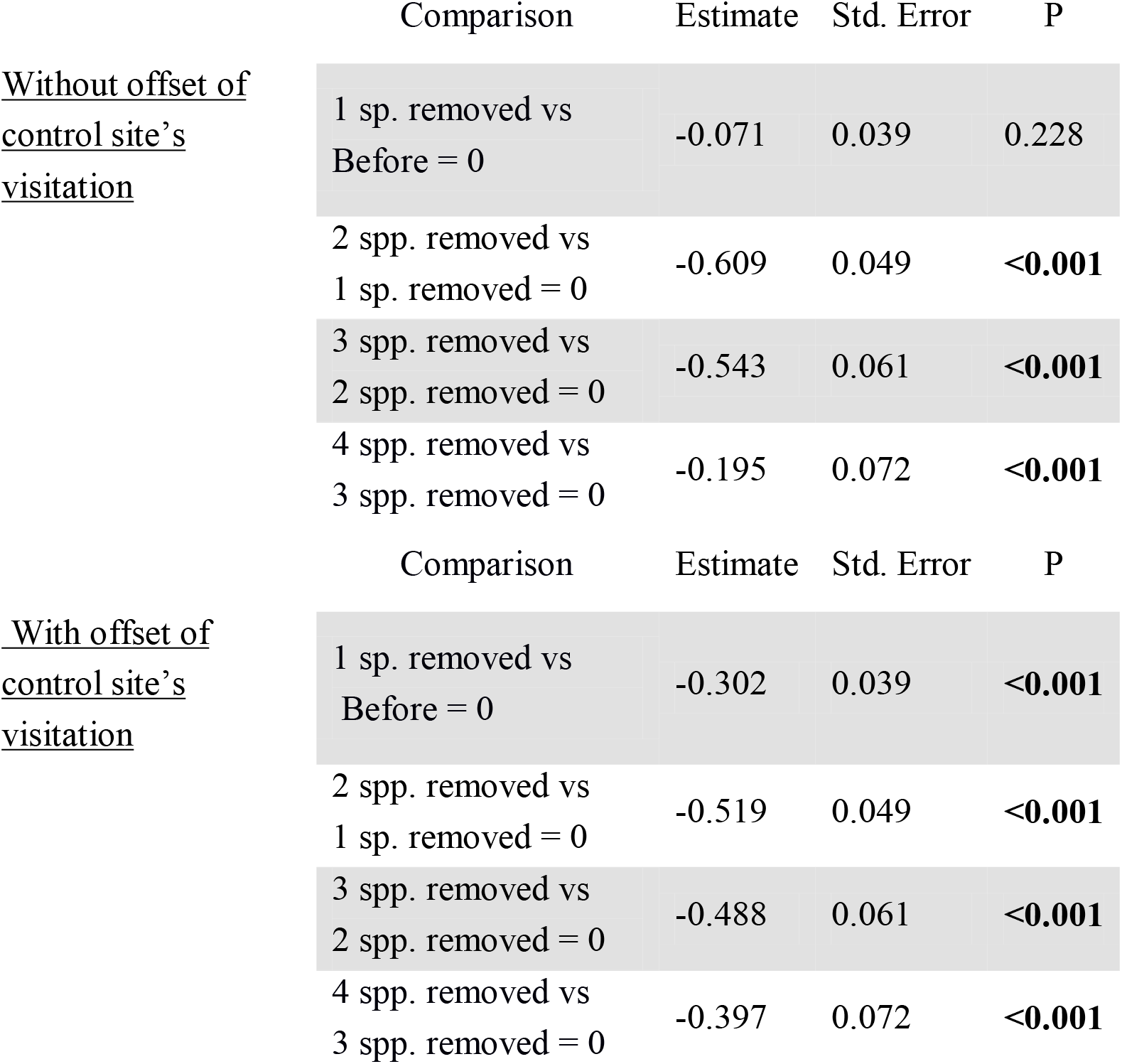
Multiple comparison statistics on the effect of treatment on general visitation, i.e. the total abundance of flower-visiting insects per transect, in the sequential removal experiment, with and without an offset with the visitation in the control site. The treatment of sequentially removing four most visited plant species is coded as “Before”, “1 sp. removed”, “2 spp. removed”, “3 spp. removed”, “4 spp. removed”. Statistically significant effects (P<0.05) are highlighted in bold.

|  | Comparison | Estimate | Std. Error | P |
| --- | --- | --- | --- | --- |
| <u>Without offset of control site's visitation</u> | 1 sp. removed vs Before = 0 | -0.071 | 0.039 | 0.228 |
|  | 2 spp. removed vs 1 sp. removed = 0 | -0.609 | 0.049 | **<0.001** |
|  | 3 spp. removed vs 2 spp. removed = 0 | -0.543 | 0.061 | **<0.001** |
|  | 4 spp. removed vs 3 spp. removed = 0 | -0.195 | 0.072 | **<0.001** |
|  | Comparison | Estimate | Std. Error | P |
| <u>With offset of control site's visitation</u> | 1 sp. removed vs Before = 0 | -0.302 | 0.039 | **<0.001** |
|  | 2 spp. removed vs 1 sp. removed = 0 | -0.519 | 0.049 | **<0.001** |
|  | 3 spp. removed vs 2 spp. removed = 0 | -0.488 | 0.061 | **<0.001** |
|  | 4 spp. removed vs 3 spp. removed = 0 | -0.397 | 0.072 | **<0.001** |

### Visitation of individual plant species

Visitation at the level of individual plant species was highly variable and significantly dependent on the number of plant species removed during the experiment (**χ**^2^= 11.39, df=4, p < 0.05). The average trend in plant species-level visitation was non-linear across treatment levels and differed between sites. (Fig. 2).

Plants showed high variation in visitation both within sites (among transects) and between sites (coloured lines and confidence intervals in Fig. 2). Nevertheless, the trend was not uniform across plant species, as idiosyncratic responses took place: visitation to some plants increased while in others it decreased in response to the same level of the removal treatment.

**Figure 2.**
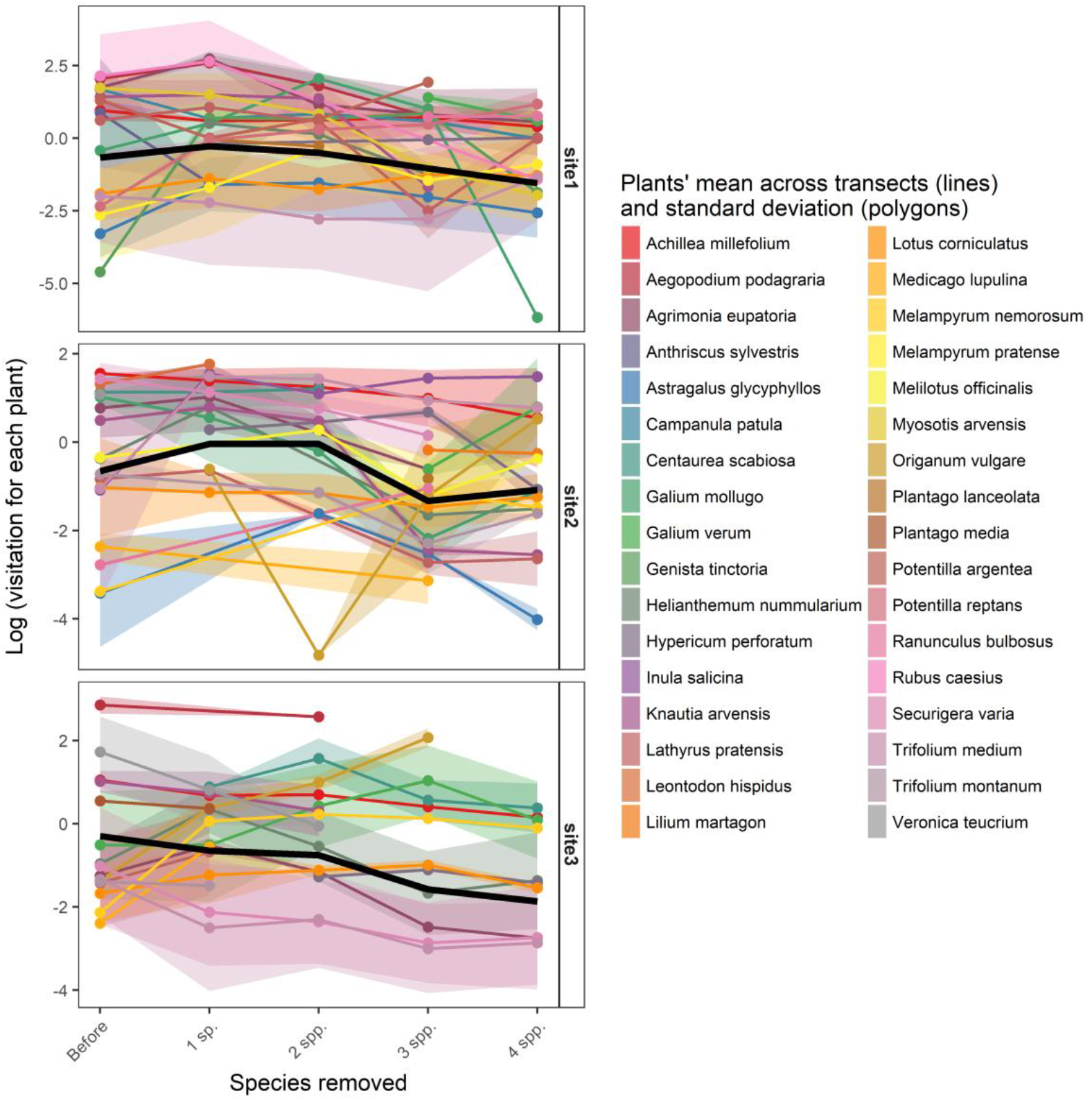
Trends of flower visitor abundances (“visitation”) on individual plant species in the sequential removal experiment. Visitation is shown on a logarithmic scale, coloured lines are estimated means across transects of a given plant species, coloured polygons are standard deviation around the plants’ means obtained to indicate variation among transects, black line is the plant community average trend.

### Plant traits

All plant-traits included in the models except colour and plant height were significant predictors of insect visitation (Table 3). Our results show that after the removal of a plant, the visitation remained stable in plants with the same flower shape, but decreased in plants with flower shapes different to the removed species. Furthermore, visitation increased in plants with relatively small inflorescences and high sugar content per flower (Fig. 3).

**Table 3.** The effect of plant traits on changes of visitation after plant removal evaluated by single-term deletion of interaction terms from a GLMM model. Columns are used for interactions between treatment (“Treatment” referring to before-after removal of one plant species) and difference of trait values of individual plants from the removed species. Trait difference for colour and flower shape was classified as “same” or “different”. Log-ratio difference was used for plant height (“Height”), inflorescence size (“Size”) and amount of sugars in nectar (“Sugars”). ΔAIC refers to the change of AIC after removing the tested term from the full model. Statistically significant effects (P<0.05) are highlighted in bold.

| | ΔAIC | $\chi^2$ | <u>P</u> |
| --- | --- | --- | --- |
| Treatment : Color | 0.4 | 2.441 | 0.628 |
| Treatment : Shape | 14.8 | 16.845 | **<0.001** |
| Treatment : Size | 16.5 | 18.573 | **<0.001** |
| Treatment : Height | -2.0 | 0.027 | 0.870 |
| Treatment : Sugars | 3.3 | 5.359 | **0.021** |

**Figure 3.**
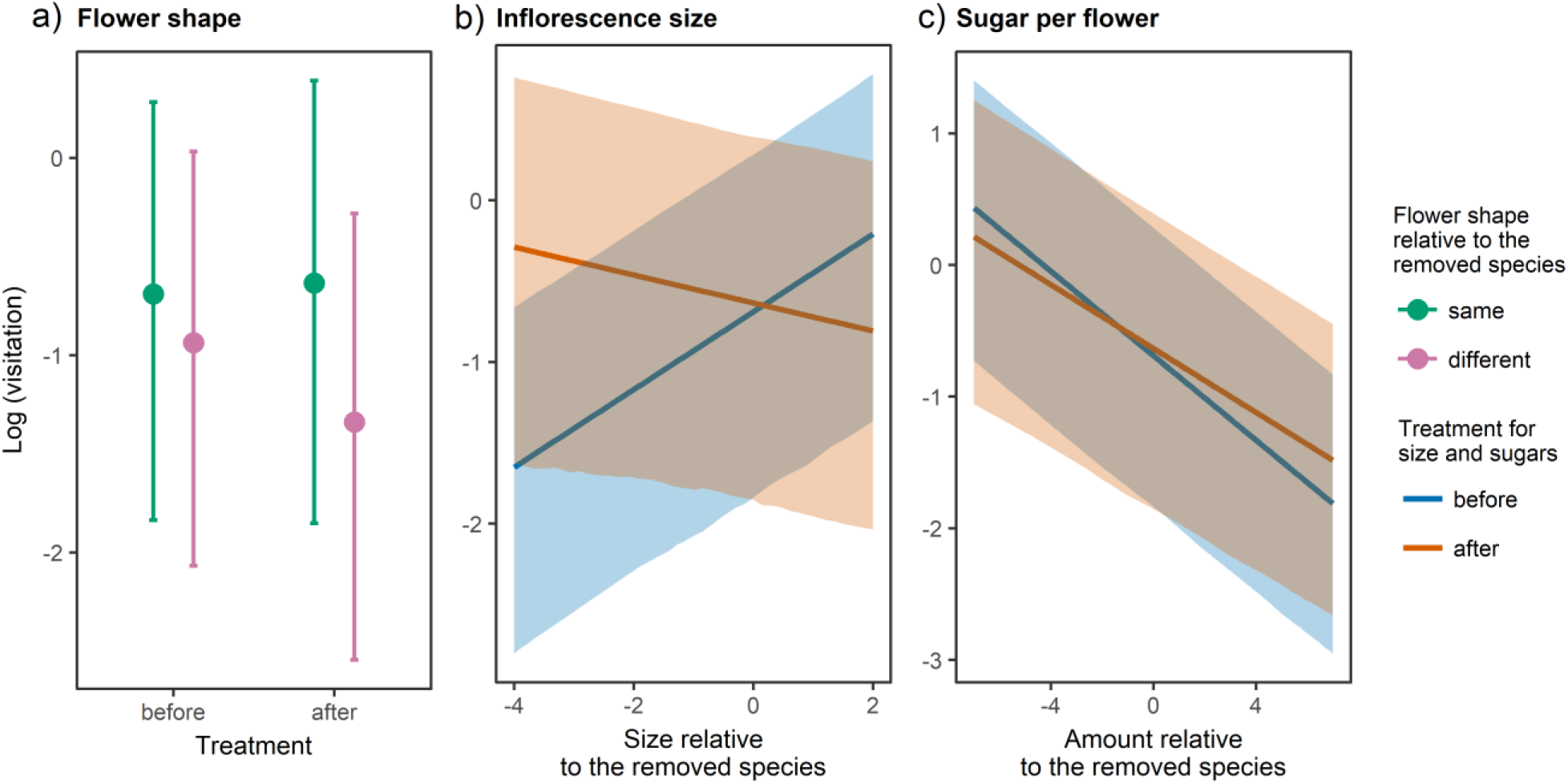
Sequential removal experiment’s flower visitor abundances as a response to plant functional traits expressed as the difference between a given plant trait value and the trait value of the removed plant species across treatment. Estimated (modelled) means and confidence intervals are represented. Only the statistically significant traits are presented (plot a-c).

### Pollination effectiveness

The number of pollen tubes per stigma fluctuated significantly during the experiment, but there was no consistent trend (Fig. 4). Treatment was a significant predictor of pollen tube number (**χ**^2^= 19.9, df=4, p < 0.001), but visitation was not (**χ**^2^= 0.62, df=1, p = 0.43).

**Figure 4.**
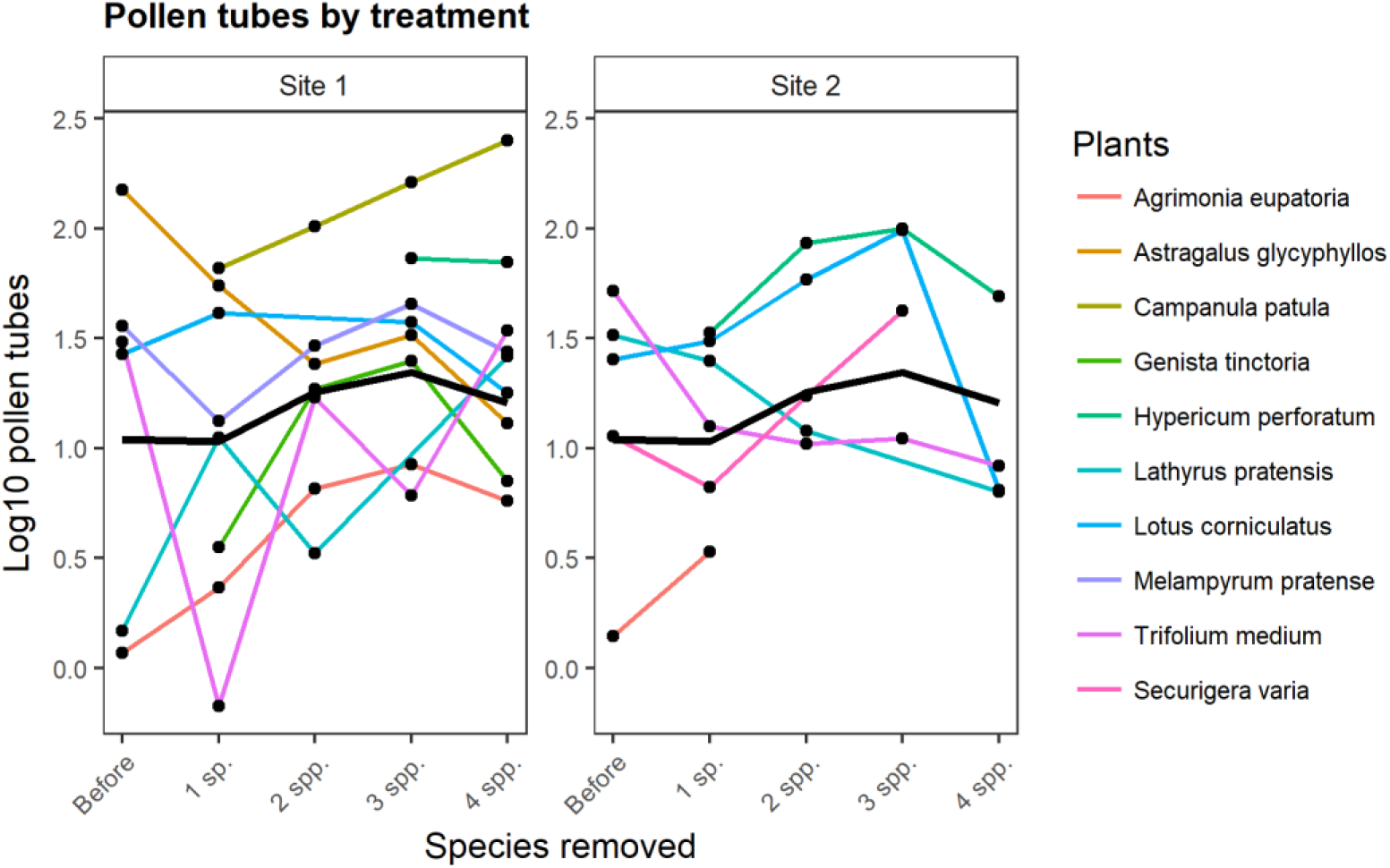
Trends of the number of pollen tubes in two of the sites in the sequential removal experiment. The number of pollen tubes is shown on a logarithmic scale, coloured lines are the estimated means across transects of a given plant species, the black line is the plant community average trend.

The average trend of pollen tube number was not linear but fluctuating (Fig. 4), because it decreased after the first species had been removed but it slightly increased during the following removals, then decreased again after the last species had been removed. However, the trend was not uniform across plant species, as idiosyncratic responses took place (coloured lines in Fig. 4).

### Standing crop of nectar

The standing crop of nectar did not change significantly as plants were removed (Fig. 5) (**χ**^2^= 2.14, df=4, p = 0.71) and visitation had no significant effect (**χ**^2^= 0.12, df=1, p = 0.71).

**Figure 5.**
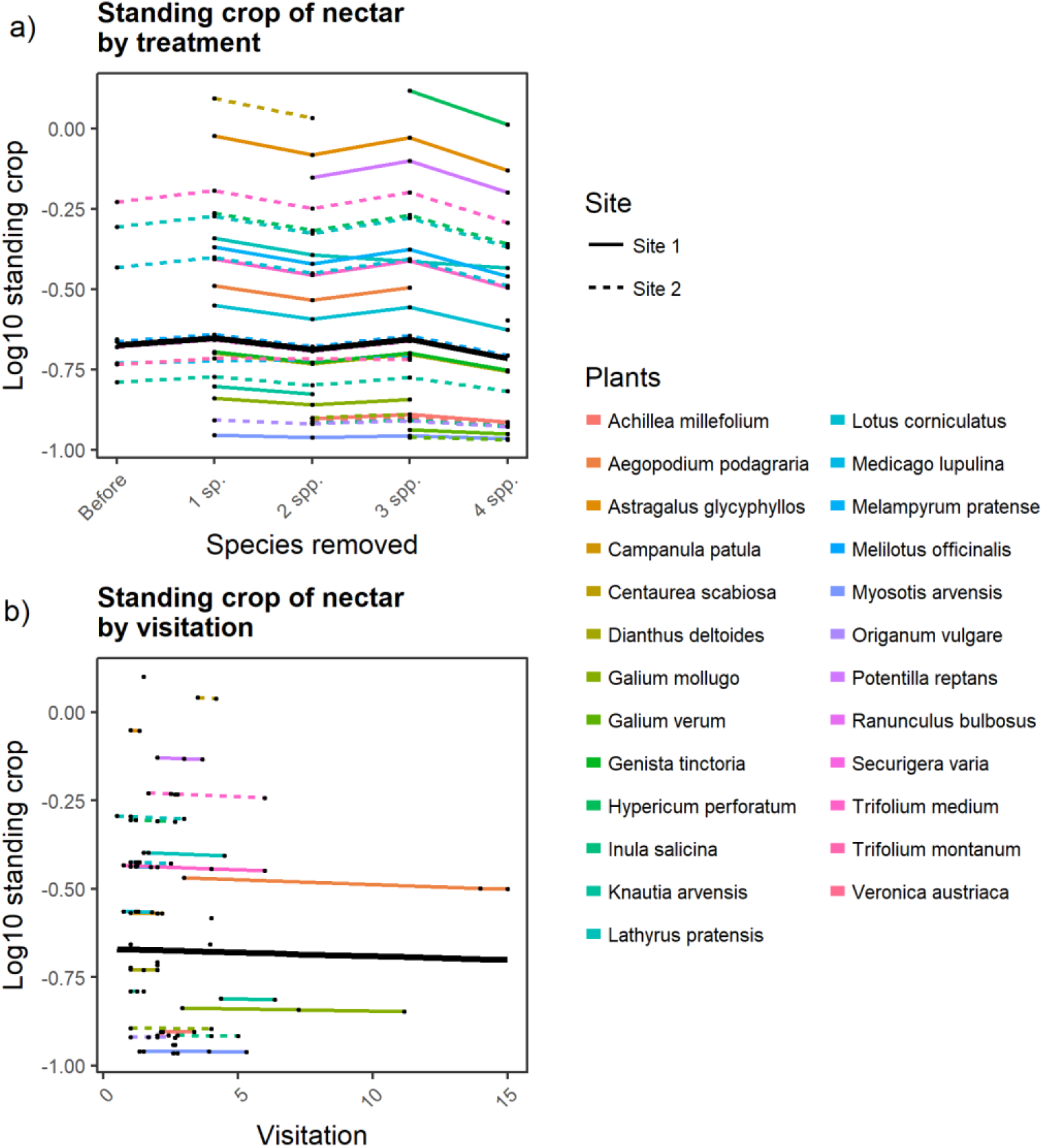
Standing crop in both sites in logarithmic scale vs. Treatment (plot a) and flower visitor’s visitation (plot b), coloured lines are the estimated means across transects of a given plant species, black line is the community-average trend.

## Discussion

We demonstrated that removal of several generalised plant species led to changes in overall flower visitation at the level of the entire community, but that visitation and pollination of individual plant species were affected in a species-specific way. We hypothesised that, at the community level, flower visiting insects may respond to the removal of the most visited plants according to one of three scenarios: (a) they may redistribute their visits equally among the remaining plants, (b) they may switch to a subset of the remaining plants depending on the traits of the plants, or (c) they may stop foraging at the affected sites. Our results strongly support the scenario (b) and partly also (c).

In the pilot study, total visitation did not change after the removal of the most visited plant. Flower visitors responded to the removal of their main food source by increasing their visitation to the next most visited plant, while visitation of the remaining plants was unaffected. The observed shift of pollinators between the two most important species happened because the second generalist plant assumed the role of the removed generalist plant^20^. There was no sign of insect emigration in the pilot study after removing one dominant food source, because the nearby control site changed only slightly and the visitation did not increase on the plant species that was removed in the treated site. More pronounced changes occurred in the sequential removal experiment, where 4 plant species were removed, one at a time. In this latter experiment, pollinators distributed preferentially on a subset of the plants remaining in the experimental site after removal (scenario “b”), instead of visiting all other plants (scenario “a”) or being unable to use alternative resources (scenario “c”).

The main outcome is that some pollinators shifted between plant species but their resource use depended on how plant functional traits related to the removed plants’ traits. Data from both the pilot and sequential removal experiment thus support a conclusion that flower visiting insects react to the loss of important resources by selectively increasing their visitation to a subset of the remaining plants (scenario “b”).

Scenario “b” has implications for the entire plant community and reveals a key feature of complex interacting communities. That is, a few dominant, generalist species support the overall interactions structure by facilitating many other species, as outlined by ^34^. This community-level facilitation likely took place also in the treated sites of our experiments, as the presence of a small number of most generalist plants supported high visitation in the plant assemblage. This reinforces the hypothesis that complex community-level interactions are based on a core of a few important species, without which the system appears as altered and impoverished ^12^. This pattern has important implications for both conservation of community-level interactions and functionality of these systems, as we discussed at the end.

Removal of multiple generalist plant species in the sequential removal experiment led to a decrease of visitation both at the site level (total visitation per transect) and on average also at the single plant species level. This might suggest that some pollinators did not find alternative resources as the removal of plants continued and insects stopped foraging at the sites. We did not collect any data on insect dispersal, so we do not know whether emigration increased after the removal. However, other aspects suggest that the remaining impoverished plant community was not affected by insect emigration. Specifically, pollination effectiveness fluctuated; the nectar consumption did not change as indicated by a nearly constant nectar standing crop; instead, both pollination effectiveness and nectar consumption were expected to decrease in scenario “c” as a consequence of a drop in local pollinator visitation.

Although previous simulation models of the consequences of species extinctions in plant-pollinator networks did not account for the redistribution of pollinators following the removal of key resources ^23,35^, our study shows that static responses in foragers are not the rule. Instead, these assemblages are dynamic and new interactions can be established after perturbations ^17,24^. Adaptive foraging by consumers within a food web has been suggested to be important for the stability of complex communities^36^. This is because perturbations to the system are buffered by switches in interactions and use of new resources^24^. However, in our results, utilization of new resources investigated at the level of the entire pollinator guild was not random but was constrained by particular plant traits. Specifically, the flower visitors did not swap between flower shapes after removals of plants, and thus the flexible foraging of insects was constrained by flower morphology. The sugar content of nectar also affected the flower visitors because flower visitors were attracted by more rewarding flowers once the plant community was impoverished. Thus the energetic intake also constrained the process of utilizing new resources. Furthermore, smaller inflorescences also became more visited after removals. It is likely that they were underutilised in the original community because of their small floral display, so their visitation benefited from the overall reduction of flower abundance after the removals. Smaller inflorescences may also provide more resources per flower, and thus be more rewarding to flower visitors on a per-visit basis^38^. Taken together, our results imply that foraging can be flexible but also constrained within a specific plant-trait space^39^. These constraints would eventually limit the accessibility of new resources after perturbations.

Another aspect of the complexity that emerged after the experiments is that pollinator service varied across the treatment in a non-linear way, as shown by the fluctuating pattern of pollen tube numbers. This might be due to destabilization of the pollinators by the removal of key resources, which thus reacted in a fluctuating way. In fact, exclusion experiments have highlighted that in the absence of a dominant pollinator, other pollinators can compensate by becoming themselves more effective pollen vectors^22^. However, another work has shown that once the abundant pollinator is excluded, plant fitness and fidelity to flowers can decrease^21^. Although these two studies contradict each other, our results showed both responses: increasing or decreasing pollination effectiveness in different plant species. This resulted from species-specific responses as some plants benefited by receiving more pollen, while other species received less, after removing key plant species. Thus, an idiosyncratic, fluctuating trend of pollination was the outcome at increasing impoverishment of the plant assemblage.

Further variation was found when considering the responses in visitation on each plant species; some plant species had different trends compared to the average community trend after treatment (Fig. 2). Such plants could be highly competitive taxa, able to gain pollinator visitation above average while other plants were losing visitors. From this fact, it is possible to indirectly draw another conclusion about the most generalist plants. That is that they facilitate only a subset of species by keeping visitation on that subset high, while other plants are being limited and receiving less visitation. Such species-specific facilitation of pollination has been observed also in the case of invasive species (e.g.^19^). This could also be responsible for high spatial variation in pollinator abundances within sites (see the wide standard deviation around plant means in Fig. 2, which reflects differences between transects). That is, a given plant species had high visitation in one sampling transect and low in another transect, even on the same day. Although these patchy responses could be due to a local heterogeneity of abiotic factors^46^, it seems more likely that the set of neighbouring plant species and their relative abundances caused variations in competition and facilitation^44^. In other words, in one patch the facilitation might predominate, while competition might outweigh facilitation in another patch, thus resulting in very complex overall patterns^6^.

The consequences of species loss observed in our experiments have several implications for conservation of biodiversity. Generalist plants which are visited by a wide range of insects play a key role in plant-flower visitor interactions, and they serve as hubs in the interaction networks ^5,20,50^. The removal of such generalist plants in our experiment led to important decreases of visitation and significant fluctuations of pollination effectiveness. We also showed that some of the responses were highly variable and species-specific. Furthermore, the insects changed their use of resources after the perturbation but their foraging flexibility was constrained by plant traits, which likely limited the utilization of new resources after plant removal. Thus, the stability of this system could depend on a small subset of important species whose loss has severe consequences for the entire community of plants and flower-visiting insects. We thus conclude that generalist plants play a key role in sustaining the complex pattern of interactions in the community^5,12^ and may be more important than commonly thought for the conservation of species-rich ecosystems^51,52^.

## Acknowledgements

PB, JK and AA would like to thank Dagmar Hucková, Tomáš Gregor, Michal Rindoš, Michal Bartoš, and Zuzana Chlumská for their help during field work in CZ. In the UK, JO & ST wish to thank The Wildlife Trust for Bedfordshire, Cambridgeshire, Northamptonshire and Peterborough for permission to work at Bradlaugh Fields, and the field assistants Joanna Tarrant, Lee Muttock, Claire Hepinstall Md Lutfor Rahman, and Christine Berrill. This project was supported by the Czech Science Foundation (projects GP14-10035P and GJ17-24795Y) and AA and PB were also supported by a grant GA JU 152/2016/P provided by the University of South Bohemia. Funding to JO and ST was provided by The Dr Mike Daniel Bursary of The University of Northampton. The funders had no role in study design, data collection and analysis, decision to publish, or preparation of the manuscript.

## Supplementary material

**Summery material Figure S1.**
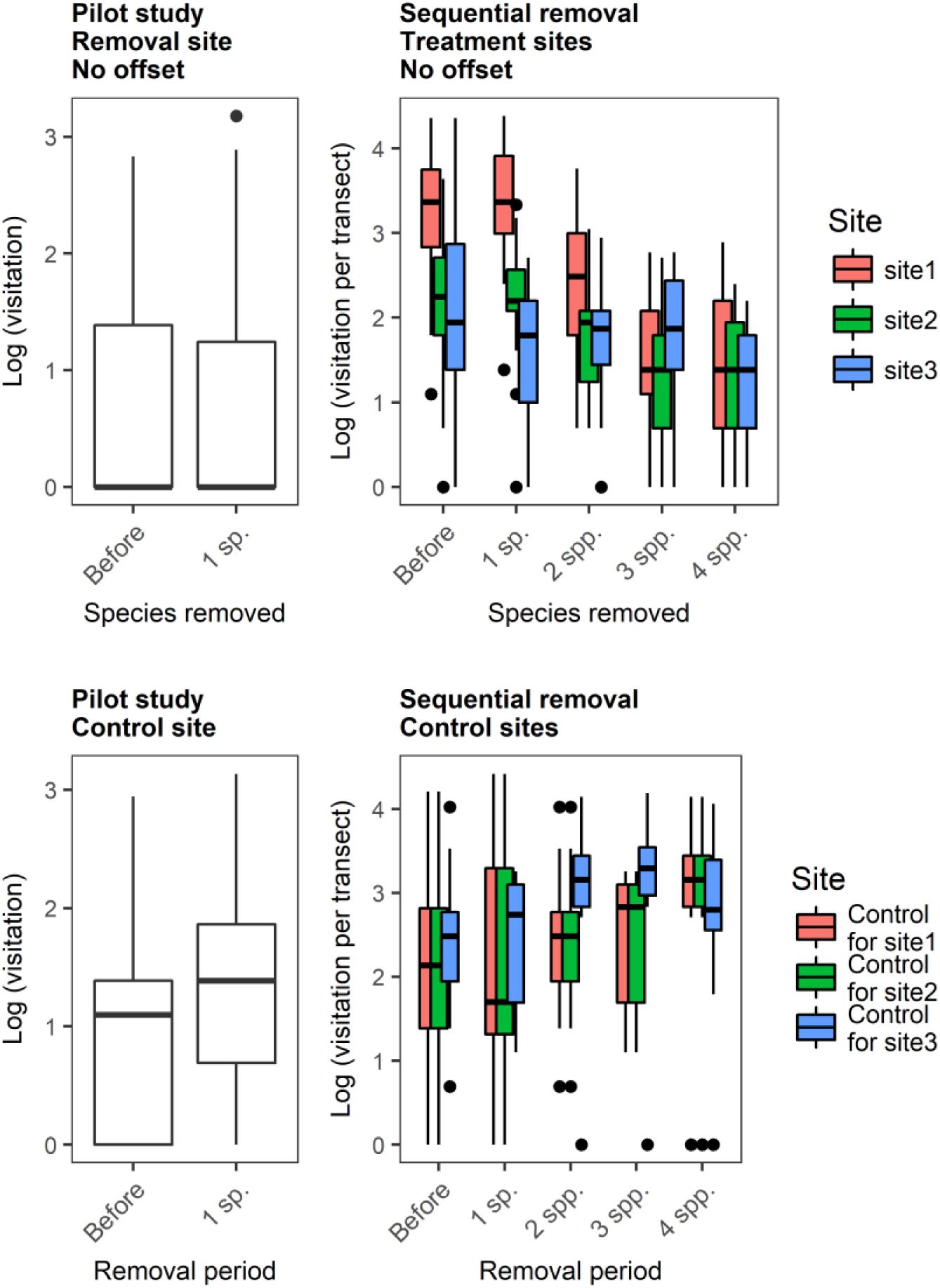
Overall visitation in control sites of the pilot and sequential removal experiment.

